# MCPNet : A parallel maximum capacity-based genome-scale gene network construction framework

**DOI:** 10.1101/2022.07.19.500603

**Authors:** Tony Pan, Sriram P Chockalingam, Maneesha Aluru, Srinivas Aluru

## Abstract

**Motivation:** Gene regulatory network (GRN) reconstruction from gene expression profiles is a compute- and data-intensive problem. Numerous methods based on diverse approaches including mutual information, random forests, Bayesian networks, correlation measures, as well as their transforms and filters such as data processing inequality, have been proposed. However, an effective GRN reconstruction method that performs well in all three aspects of computational efficiency, data size scalability, and output quality remains elusive. Simple techniques such as Pearson correlation are fast to compute but ignore indirect interactions, while more robust methods such as Bayesian networks are prohibitively time consuming to apply to tens of thousands of genes.

**Results:** We developed MCP Score, a novel maximum-capacity-path based metric to quantify the relative strengths of direct and indirect gene-gene interactions. We further present MCPNet, an efficient, parallelized GRN reconstruction software based on MCP Score, to reconstruct networks in unsupervised and semi-supervised manners. Using synthetic and real *S. cervisiae* datasets as well as real *A. thaliana* datasets, we demonstrate that MCPNet produces better quality networks as measured by AUPR, is significantly faster than all other GRN inference software, and also scales well to tens of thousands of genes and hundreds of CPU cores. Thus, MCPNet represents a new GRN inferencing tool that simultaneously achieves quality, performance, and scalability requirements.

**Availability:** Source code freely available for download at *https://doi.org/10.5281/zenodo.6499748* and *https://github.com/AluruLab/MCPNet*, implemented in C++ and supported on Linux.

**Supplementary information:** Supplementary data are available at *Bioinformatics* online.

## 1 Introduction

Gene regulatory network (GRN) inference from high-throughput gene expression data is a compute intensive task. Although several different inference algorithms based on Bayesian networks (Hartemink, 2005), mutual information (MI) theory (Faith *et al*., 2007; Aluru *et al*., 2013) Pearson correlation (Langfelder and Horvath, 2008), regression (Bonneau *et al*., 2006; Huynh-Thu *et al*., 2010), and random forest (Aibar *et al*., 2017) have been developed over the past two decades, scalability remains a critical challenge when working with tens of thousands of genes and/or observations. Aluru *et al*. (2021) found that 9 of the 15 methods analyzed failed to infer GRNs from a dataset with approximately 18, 000 genes. Furthermore, of the 6 that succeeded, 4 required between 4 and 49 days to complete. To construct large GRNs with tens of thousands of genes in reasonable time, scalable algorithms and efficient parallel implementations are necessary. Arboreto (Moerman *et al*., 2019) recently introduced distributed computing capability to construct random forest-based transcription factor (TF)–target networks; however, this is still not scalable for large networks. TINGe (Zola *et al*., 2010), using a parallel MI-based approach can construct large genome-scale GRNs in a significantly shorter amount of time.

Past surveys conducted with established benchmark data suggest that Bayesian and MI–based methods are among the best performing (Marbach *et al*., 2012; Lachmann *et al*., 2016; Aluru *et al*., 2021) with respect to the quality and accuracy of inferred interactions. However, unlike Bayesian networks, MI-based methods are more amenable to large scale GRN construction (Chockalingam *et al*., 2017). One key caveat though is that these methods require post processing such as Stouffer Transform in CLR (Faith *et al*., 2007) or MI value filtering based on p-values and data processing inequality (DPI) measures in ARACNe–AP (Lachmann *et al*., 2016) and TINGe. Such filtering complicates network evaluation by standard metrics such as the area under the precision-recall curve (AUPR) as it introduces discontinuities in the value range. Additionally, DPI requires a user-supplied tolerance parameter that is challenging to determine *a priori*. As DPI reflects the information transmission capacity, it is a special case of the Maximum Capacity Path (MCP) problem in graph theory, which has previously found applications in network routing (Pollack, 1960), image compositing (Fernandez *et al*., 1998), and metabolic pathway analysis (Ullah *et al*., 2009). Formulated as such, the tolerance parameter is no longer required.

In this paper, we present a novel GRN inference approach based on MCP to characterize and compares indirect gene interactions to identify significant gene-gene interactions without thresholding or user-specified parameters. Our unsupervised approach examines all indirect paths between two genes to compute the maximum indirect interaction capacity, and allows efficient investigation of paths of arbitrary lengths. Using the same framework, we also established a semi-supervised approach to identify optimal weight for combining interactions with multiple path lengths. We further created an efficient parallel implementation of the algorithm, called MCPNet, that can scale to hundreds of cores. We evaluated performance of our unsupervised and semi-supervised approaches using MI as the direct interaction measure and paths with length up to 4, and demonstrate that our method delivers higher network quality and superior computational performance when compared to other state-of-the-art GRN inference software.

## 2 Materials and Methods

A gene expression dataset with samples *S* is formulated as a matrix *P* with |*V*| rows and |*S*| columns, each row corresponding to expression levels for a gene, and each column corresponding to the gene expression profile of a sample in *S*. A gene regulatory network is defined as an undirected weighted graph *G* =< *V, E, W* >, where *V* = {*v*_1_ … *v*_*N*_} is the set of *N* genes, *E* = {(*v*_*i*_, *v*_*j*_)|*v*_*i*_, *v*_*j*_ ∈ *V*} is the set of edges representing interacting genes, and *W* is a matrix of size *N × N*, where the entry *W*_*ij*_ is an edge weight that quantifies the interaction strength of *v*_*i*_ with *v*_*j*_.

The significance of the edge weights depend upon the underlying mathematical model. For a correlation network, *W*_*ij*_ indicate the degree of correlation between *v*_*i*_ and *v*_*j*_, for example measured using Pearson correlation. Correlation networks are generally symmetric, i.e. *W*_*ij*_ = *W*_*ji*_, but may contain negative values. In Arboreto model (Huynh-Thu *et al*., 2010), *W*_*ij*_ indicates the degree to which the expression of gene *v*_*i*_ can predict *v*_*j*_ ‘s gene expression, thus *W* is non-negative but may not be symmetric.

In case of a mutual information (MI) network, the expression values for a gene *v*_*i*_ ∈ *V* are modeled by the random variable *X*_*i*_, samples of which are values in row *i* in *P*, denoted by ⟨*P*_*i*1_, …, *P*_*i*|*S*|_⟩. From the measured gene expression profiles ⟨*P*_*i*1_, …, *P*_*i*|*S*|_⟩ and ⟨*P*_*j*1_, …, *P*_*j*|*S*|_⟩, the interaction strength *W*_*ij*_ between genes *v*_*i*_ and *v*_*j*_ can be computed by estimating the mutual information. Since mutual information is non-negative and commutative, *W* is symmetric and positive semi-definite.

In this paper we primarily use MI to demonstrate the performance our proposed method, though it is agnostic of the semantics of the *W* matrix or its symmetry. For *W* with negative values, our proposed method can use the magnitude, i.e. absolute value, of the interaction strength instead.

### 2.1 MCP Score

Longer range interactions are common in eukaryotic organisms (Vermeirssen *et al*. (2014); Lu *et al*. (2007); Itzhack *et al*. (2013); Cowen *et al*. (2017)). In these cases, an interaction between two genes is mediated through one or more intermediary genes and can be modeled as a length-*L* path in *G*. Reconstructing the GRN from gene expression profiles is guided by two goals: identifying strong direct interactions between gene pairs, and simplifying the network by removing redundant direct and indirect interactions. Our proposed method seeks to accomplish both by first identifying the strongest indirect interactions between two genes, then compare it to the direct interaction strength. Direct interactions are kept only if they are stronger than the strongest indirect interactions.

The maximum indirect interaction strength is determined by solving the maximum capacity path (MCP) problem. Figure 1 illustrates different length-2 and length-3 paths between genes *v*_*s*_ and *v*_*t*_ as dashed lines and their direct interactions as solid lines. For each path, the minimum edge weight is its maximum capacity. Among all the possible length-*L* paths between *v*_*s*_ and *v*_*t*_ in *G*, the maximum capacity path is the path with the highest minimum edge weight. We use 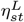 to denote the capacity of such a path between *v*_*s*_ and *v*_*t*_, and refer to it as *L-path Capacity* or *Path Capacity*. Formally,

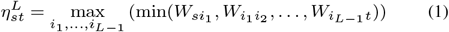

**Fig. 1:**
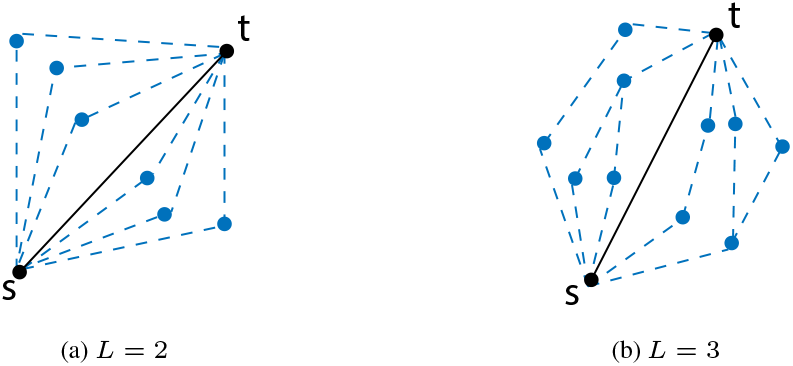
Computing 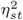 and 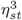 in a network requires exploring different paths with one and two intermediate vertices respectively, as shown above.

Once computed, 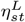 is compared to the direct interaction strength *W*_*st*_. We define the *L*-path MCP score between the genes *v*_*s*_ and *v*_*t*_, denoted as 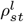, as the following ratio:

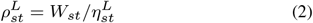

We use both the terms *L-path MCP Score* and *MCP Score* to refer to 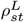. We denote ℛ^*L*^ as a matrix of all the 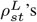 for every pair of *s* and *t*. As a ratio of the edge weight to the path capacity, we expect 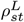 to be an indicator of the interaction strength of genes *v*_*s*_ and *v*_*t*_ while factoring in the effect of other interactions in the *L*-neighborhood of *v*_*s*_ and *v*_*t*_. We note that the DPI-based approach used by ARACNe-AP and TINGe can be modeled as a specialization of the *L*-path MCP score, as shown in Supplementary Section S1.1. For indirect interactions involving two, three or four intermediate interactions, the *η*_*st*_ definitions are:

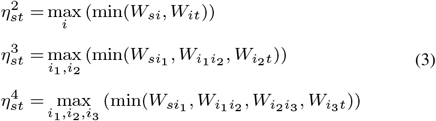

### 2.2 MCP Score Algorithm

The capacity 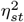 can be computed explicitly by enumerating all possible length-2 paths between *v*_*s*_ and *v*_*t*_. Algorithm 1 shows the pseudo-code for this approach. The 2-path MCP score defined in Section 2.1, 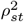, can be calculated for the complete network directly by first computing 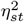 for a pair of genes via Algorithm 1, iterating over the two gene expression profiles in *O*(|*V*|) time. The algorithm has a run-time complexity of *O*(|*V*|^3^) as it iterates over all elements of a |*V*| × |*V*| matrix (Algorithm 2).

Algorithm 2 applies for *L*-path MCP scores for any arbitrary *L*. Naive extension of Algorithm 1 to longer range interactions is exponential in computational complexity, however, as an increase of *L* by 1 increases computation by |*V*|-fold. For an indirect interaction of length *L*, the naïve 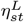 computation complexity is *O*(|*V*|^*L−*1^), leading to *O*(|*V*|^*l*+1^) to compute the scores for all gene-gene pairs.

Pollack Pollack (1960) showed that the maximum capacity 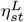 of all length-*L* paths between two vertices in a graph can be computed efficiently via recursive path bisection. Formally,

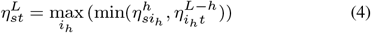

where *h* = ⌊*L/*2⌋. The length-2 realization, 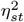, in equation 3 forms the base case for the recursion. This partitioning leverages the associative properties of the maximum and minimum operations. Let 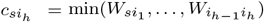 and 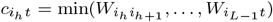.

Equation 1 can then be rewritten as

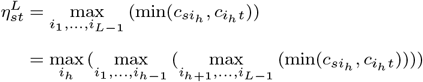

As 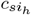 does not depend on *i*_*h*+1_ … *i*_*L−*1_, it is effectively a constant with respect to the 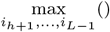 operation. Since the maximum and minimum operators are distributive over each other with respect to a constant, *i.e*., max(min(*z, x*), min(*z, y*)) = min(*z*, max(*x, y*)),

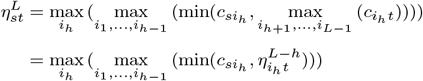

Similarly 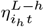 is constant with respect to *i*_1_ … *i*_*h−*1_. Applying the identity again leads to Equation 4.

#### Algorithm 1: Path Capacity Kernel for Path Length 2

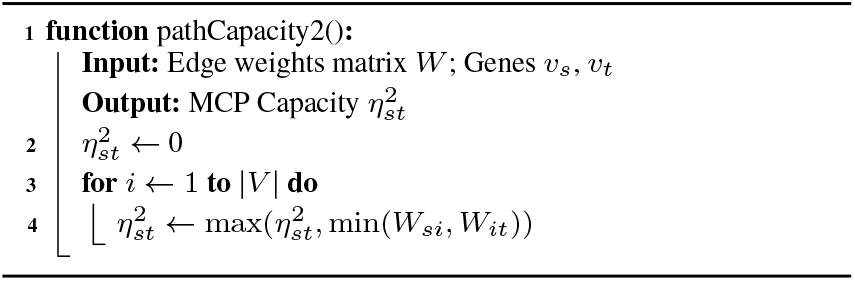

#### Algorithm 2: MCP Score

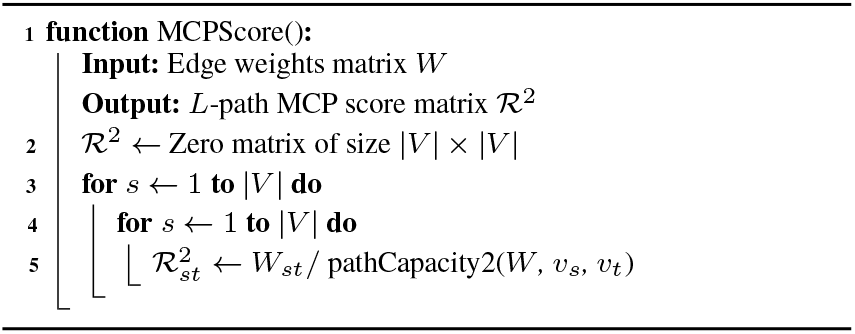

The recursive path bisection allows *L*-length MCP scores to be computed in *O*(|*V*| log_2_ *L*) for a single gene-gene pair, and the long range DPI scores for all gene pairs to be computed in *O*(|*V*|^3^ log_2_ *L*) time. A realization of Equation 4 is shown in Algorithm 3. For a gene pair (*v*_*s*_, *v*_*t*_), the *η* values for half-length long range interactions between *v*_*s*_ and *v*_*t*_ with all possible intermediary genes *v*_*r*_ ∈ *V* are computed. The *η* value for (*v*_*s*_, *v*_*t*_) is then computed following the same maximum capacity path problem solution.

### 2.3 Parallel Implementation

MCPNet pipeline consists of a sequence of functions. We first compute the mutual information (MI) between pairs of genes, using the gene expression profile matrix as input. The resulting MI matrix was then transformed to MCP score via the algorithm described in the sections above. Our implementations of algorithms 1, 2, and 3 are optimized and parallelized for multi-core and multi-node systems.

First, we note that Algorithm 2 has data access pattern identical to matrix-matrix multiplication, with Algorithm 1 replacing vector dot product as the computational kernel. Indeed, fast matrix multiply algorithms have been applied to the maximum capacity path problem (Vassilevska *et al*., 2007; Duan and Pettie, 2009) to achieve sub-cubic run-time. For simplicity and ease of parallelization, we compute each *η* directly in *O*(|*V*|) time. In contrast to the *dominant product* algorithm by Vassilevska *et al*. (2007) that requires random memory access, our algorithm incurs only sequential memory access and therefore is cache-friendly and can leverage hardware memory prefetching.

Since each element can be computed independently from the other elements of the output matrix, a simple parallelization scheme is sufficient. The output matrix is partitioned into 8 × 8 tiles and tiles are assigned to the compute nodes and CPUs. Use of tiles improves cache reuse as the 8 rows and 8 columns of the input matrices needed to compute one tile are likely to remain in cache memory during computation.

To ensure that the required input matrix row and column to compute one 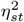 value are in local memory, the mutual information matrix is replicated on each compute node, and shared between cores of the same node. The MI computation is similarly organized across all nodes, with the gene expression profile matrix replicated and the MI matrix tiles partitioned across cores. One round of all-to-all personalized communication via Message Passing Interface (MPI) reconstructs the MI or MCP score matrices on all compute nodes. The input replication reduces communication during computation. The overall parallelization approach is shown in Figure 2.

**Fig. 2:**
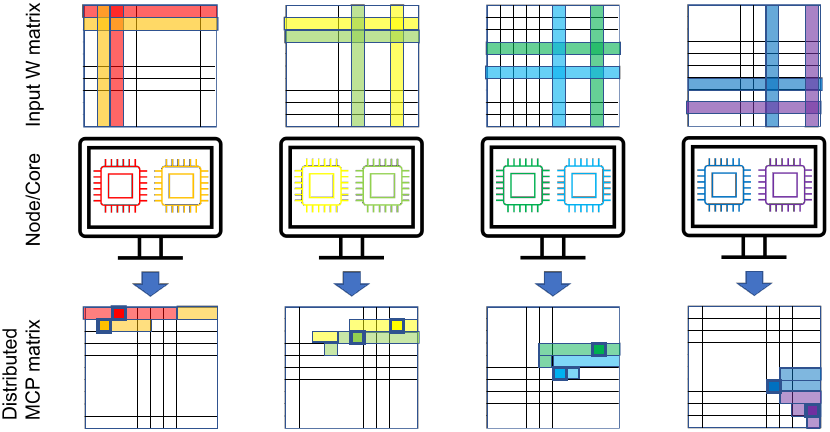
Paralellization scheme for the MCP Score algorithm. The input W matrix is replicated to all nodes and shared between cores of a node. Each core (colored) computes some tiles (light shading in the MCP matrix) in the MCP matrix thus the output matrix is distributed across the compute nodes. For each MCP tile being computed (solid square with thick border in the MCP matrix), the required rows and columns are shaded in the W matrix according to the core color.

#### Algorithm 3: Recursive Path Capacity Kernel for Length *L*

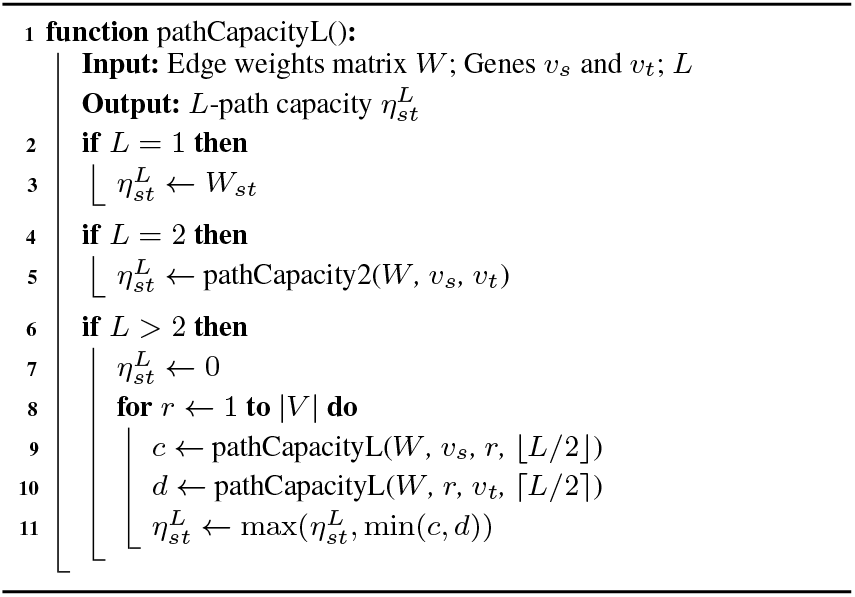

Finally, the computation to produce 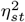 as depicted in Equation 3 is readily vectorized using element-wise minimum and maximum operators. For MI as the gene-gene interaction strength measure, since *W* is symmetric, the MCP score matrix is symmetric as well thus half of the computation can be avoided.

### 2.4 Semi-supervised Ensemble Multipath MCP scores

MCP scores as defined in Section 2.1 are for a fixed path length. To account for indirect interactions of multiple lengths, we consider a linear combination of the *η* values in computing an ensemble DPI score:

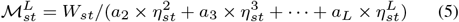

where *a*_2_, *a*_3_, …, *a*_*L*_ are coefficients for the different indirect interaction path lengths, with the standard constraint that the weights add to 1.0.

The optimal hyper-parameters are generally challenging to determine. We propose to use a semi-supervised approach to help optimize the hyper-parameters. For large whole genome regulatory networks, there are known and experimentally validated gene-gene interactions. Using the known interactions as partial ground truth for optimizing the hyper-parameters, the proposed semi-supervised approach promises to improve predictions of unknown interactions in the remainder of the network.

For the ensemble multi-path MCP score, we compute a weighted average of the 2-, 3-, and 4-path capacities by iterating over combinations of coefficients at a default interval of 0.1 for each coefficient. The AUPR of the resulting networks are computed based on known interactions. The coefficient combination with the maximum AUPR is used as the optimal combination for the ensemble. The semi-supervised approach allows the network inferencing process to adjust to the dataset available, the organism of interest, and the available ground truth.

## 3 Evaluation Methodology

We evaluate performance of MCPNet using both simulated and real datasets, and compare it with seven other popular GRN inference software – four MI-based methods (ARACNe-AP (Lachmann *et al*., 2016), CLR (Faith *et al*., 2007), MRNET(Meyer *et al*., 2008), TINGe (Aluru *et al*., 2013)), a Pearson correlation-based method, (WGCNA (Langfelder and Horvath, 2008)), a random forest-based method (Arboreto (Aibar *et al*., 2017)), and a regression-based method (Inferelator (Bonneau *et al*., 2006)). Arboreto and Inferelator are ensemble methods, in contrast to the other methods referenced above which are standalone. Each method evaluated is executed with its default parameters. We applied the standard statistical measure AUPR for assessing inferred networks. AUPR is the area under the Precision-Recall curve, and is recommended in cases where there is an imbalance of positive and negative edges in the underlying network (Davis and Goadrich, 2006). For *in silico* evaluations, we used NetBenchmark (Bellot *et al*., 2015), an R package to generate yeast simulated datasets from 2000 observations and 2000 genes. Noise are then injected at five different levels in ten replicas to create 51 total datasets.

To determine accuracy and scalability of the GRNs with real-world data, we used gene expression data from two different organisms - *S. cerevisiae* and *A. thaliana*. The *S. cerevisiae* (yeast) dataset is a compilation of multiple yeast RNA-seq expression studies, and contains 2, 577 observations and 5, 716 genes (Tchourine *et al*., 2018). To determine the quality of yeast GRNs, we utilized a set of true positives and true negatives from Castro *et al*. (2019) as ground truths. The *A. thaliana* microarray data was downloaded from public repositories (Aluru *et al*., 2021), and processed according to (Chockalingam *et al*., 2016). Data from six different tissues/conditions with varying sizes of gene expression datasets (Table 1) were used for GRN inference.

**Table 1:** *A. thaliana* microarray datasets. The number of CEL files and genes remaining after data normalization and filtering are as given.

| Tissue/Condition | No. CEL files | No. Genes |
| --- | --- | --- |
| Seed | 787 | 19120 |
| Flower | 920 | 17608 |
| Hormone | 1708 | 17753 |
| Seedling (1wk) | 2822 | 16298 |
| Leaf | 3564 | 16887 |
| Development | 5102 | 18373 |

*A. thaliana* network(s) were evaluated using interactions from the following two known networks as ground truths: (1) Arabidopsis Transcriptional Regulatory Map (ATRM) constructed from *a priori* biological knowledge by Jin *et al*. (2015). (2) N-response DFG network is a network constructed using dynamic factor graphs with time-series data from Nitrogen-treatment experiments (Brooks *et al*., 2019). The interactions from these two given networks can be considered as positives *i.e*., interactions that are expected to be highly weighted edges in any predicted network. For true negatives, we used a set of 4, 347 interaction pairs between chloroplast-encoded (CG) and mitochondria-encoded (MG) genes (Aluru *et al*., 2021).

All software were run on a computing cluster with each node having two 2.7 GHz 12-Core Intel Xeon Gold 6226 Processor processors and 256 GB of main memory and running RedHat Enterprise Linux (RHEL) 7.0 operating system. For the simulated and *S. cervisiae* RNA-seq datasets, all 24 cores of a node were used for software that could be run with multiple cores. For example, Arboreto, WGCNA, TINGE, Inferelator and MCPNet are capable of using all the cores in a node. For *A. thaliana* datasets, we used up to 8 nodes and all of the 192 cores distributed across these 8 nodes for Arboreto, TINGE and MCPNet, which are capable of using multiple distributed cores. While Inferelator is also capable of utilizing multiple shared cores, it was not used in this case due to extremely long run-times. Scripts used for the evaluations presented here can be accessed at *https://doi.org/10.5281/zenodo.6499756*.

## 4 Results and Discussion

The proposed MCPNet method takes as input correlation measures, which for the purpose of evaluation presented here are the MI values between pairs of gene expression profiles. We evaluated three common methods for estimating MI values from observed data and chose Adaptive Partitioning (AP) with Rank Transform for its accuracy and speed (see Supplementary Section S1.2). We use our implementation of the AP MI estimation algorithm based on the hybrid multi-thread, multi-node parallelization approach in Section 2.3.

The unsupervised MCP score has a single parameter, path length *L*. Empirical testing with simulated Yeast data showed that the network quality reached maximum at *L* = 4 and improvements diminished beyond 4 (Supplementary Section S1.2). Subsequent evaluations of MCPNet, including the semi-supervised ensemble multipath scores, were conducted with *L* of 2, 3, and 4. The ensemble method optimizes the multipath score via a global coefficient search at a regular interval, which is a user-adjustable parameter. We used 0.1 as the search interval as it represents a good balance between search quality and computation time.

### 4.1 Evaluation using Simulated Yeast Expression Data

We begin by evaluating the effect of noise in the gene expression profile data on network inference quality. Simulated yeast datasets were generated with varying noise levels using the *NetBenchmark* package.

For each of the datasets MCPNet generates three different sets of networks: (i) *L*-path MCP score networks (*i.e*., MCP score matrices ℛ^2^, ℛ^3^ and ℛ^4^ from section 2.1); (ii) Ensemble networks constructed using semi-supervised method discussed in section 2.4 (*i.e*., ℳ^4^ matrix in Equation 5) constructed as the weighted combination of ℛ^2^, ℛ^3^ and ℛ^4^ matrices; and (iii) Stouffer-transformed ensemble network, where Stouffer’s *Z*-transform is applied to each of the ℳ^4^ matrices and the AUPR for the best combination is reported as 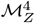. This is similar to post-MI processing in CLR (Faith *et al*., 2007) to reduce background contribution to the score.

Table 2 shows the average AUPR for networks constructed by MCPNet and other network construction methods. Results for all of the simulated data instances are given in Supplementary Table S2. The rows ℛ^2^, ℛ^3^ and ℛ^4^ are networks constructed using the path scores (Equation 2) with path lengths of 2, 3, and 4, respectively. Note that for ℳ^4^ and 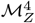 networks, the average AUPRs for the best combination is reported.

**Table 2:** Performance assessment of different GRN inference methods on simulated yeast data with noise levels ranging from 0 to 1.

| Method | 0 | 0.2 | 0.25 | 0.5 | 0.75 | 1.0 |
| --- | --- | --- | --- | --- | --- | --- |
| ARACNe-AP | 0.3503 | 0.1829 | 0.1814 | 0.0818 | 0.0349 | 0.0170 |
| CLR | 0.1523 | 0.1368 | 0.1365 | 0.1193 | 0.1035 | 0.0886 |
| MRNET | 0.2711 | 0.2257 | 0.2323 | 0.1819 | **0.1363** | **0.1075** |
| TINGe | 0.3488 | 0.2478 | 0.2482 | 0.1539 | 0.1007 | 0.0666 |
| WGCNA | 0.0549 | 0.0545 | 0.0551 | 0.0544 | 0.0523 | 0.0496 |
| $\mathcal{R}^2$ | 0.3373 | 0.1133 | 0.1157 | 0.0615 | 0.0358 | 0.0223 |
| $\mathcal{R}^3$ | 0.3690 | 0.2575 | 0.2579 | 0.1679 | 0.1029 | 0.0619 |
| $\mathcal{R}^4$ | <u>0.3938</u> | <u>0.2840</u> | <u>0.2839</u> | <u>0.1959</u> | 0.1306 | 0.0875 |
| $\mathcal{M}^4$ | **0.4198** | **0.2975** | **0.2983** | **0.2035** | <u>0.1337</u> | 0.0889 |
| $\mathcal{M}_Z^4$ | 0.2924 | 0.1380 | 0.1399 | 0.1008 | 0.0778 | 0.0576 |
| Arboreto | 0.1932 | 0.1669 | 0.1637 | 0.1349 | 0.1106 | <u>0.0902</u> |
| Inferelator | 0.2150 | 0.1849 | 0.1763 | 0.1161 | 0.0838 | 0.0638 |

Note: Numbers indicate average AUPR values over ten sampling runs. The best performing networks are highlighted in bold, and the second best is underlined. Coefficients for the best performing $\mathcal{M}^4$ and $\mathcal{M}_Z^4$ networks vary for each randomly sampled network.

As expected, the AUPR values, and hence the network performance, decreases with increasing noise levels. Our results also show that semi-supervised ensemble MCP Score achieves the best performance for low and moderately noisy datasets when compared to all the other GRN inference methods. At noise levels higher than 50%, MCPNet is the second best performing method. The selection of best coefficients for each of the *L*-path capacities (*i.e*., ℛ^2^, ℛ^3^ and ℛ^4^) in a semi-supervised manner results in an improvement over the individual MCP score networks. By exploiting known interaction information from user supplied partial ground truth, the semi-supervised approach is able to identify the best combination for each of the noise levels. Thus, MCPNet’s ensemble method is able to adjust to the unique data and noise characteristics of each dataset through fine tuning of its coefficients.

### 4.2 Evaluation with Real Gene Expression Datasets

We next evaluated performance of MCPNet using real-world data from two different organisms - yeast (*S. cervisiae*) and *A. thaliana*. For networks inferred from the real yeast data, our results show that Arboreto is the highest performing method followed by 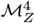 (Table 3). For this dataset, all methods showed low AUPR scores, and the absolute AUPR differences between Arboreto and 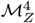 can be considered marginal. The improvement of 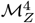 over ℳ^4^ and the relatively high AUPR for CLR suggest that Stouffer transformation as a post processing step is beneficial for this particular dataset. While Arboreto achieved the best network quality, it does so in approximately 29 hours whereas MCPNet completed the computation in 65 seconds. The quality and run-time balance strongly favors MCPNet for its ability to reverse-engineer a good-quality network in reasonable time.

**Table 3:**
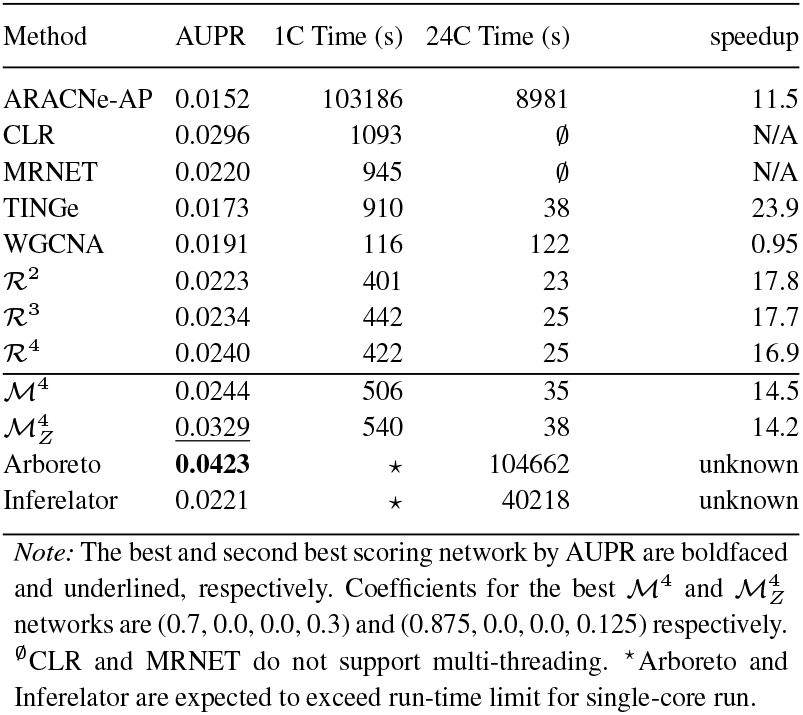
Evaluation of MCPNet’s performance with *S. cerevisiae* data. C denotes cores.

| Method | AUPR | 1C Time (s) | 24C Time (s) | speedup |
| --- | --- | --- | --- | --- |
| ARACNe-AP | 0.0152 | 103186 | 8981 | 11.5 |
| CLR | 0.0296 | 1093 | ∅ | N/A |
| MRNET | 0.0220 | 945 | ∅ | N/A |
| TINGe | 0.0173 | 910 | 38 | 23.9 |
| WGCNA | 0.0191 | 116 | 122 | 0.95 |
| $\mathcal{R}^2$ | 0.0223 | 401 | 23 | 17.8 |
| $\mathcal{R}^3$ | 0.0234 | 442 | 25 | 17.7 |
| $\mathcal{R}^4$ | 0.0240 | 422 | 25 | 16.9 |
| $\mathcal{M}^4$ | 0.0244 | 506 | 35 | 14.5 |
| $\mathcal{M}_Z^4$ | <u>0.0329</u> | 540 | 38 | 14.2 |
| Arboreto | **0.0423** | * | 104662 | unknown |
| Inferelator | 0.0221 | * | 40218 | unknown |

Note: The best and second best scoring network by AUPR are boldfaced and underlined, respectively. Coefficients for the best $\mathcal{M}^4$ and $\mathcal{M}_Z^4$ networks are (0.7, 0.0, 0.0, 0.3) and (0.875, 0.0, 0.0, 0.125) respectively. <sup>∅</sup>CLR and MRNET do not support multi-threading. \*Arboreto and Inferelator are expected to exceed run-time limit for single-core run.

*A. thaliana* networks were constructed using data from the Development category (Table 1) containing the largest number of microarray datasets. Data was processed both on one node and eight distributed nodes for all GRN methods, with the exception of Inferelator which does not support multi-node execution and whose run-time for the development dataset is expected to exceed the cluster’s job scheduler time limit. Table 4 shows that for the *A. thaliana* dataset, ℛ^4^ and ℳ^4^ produced the best results when compared to all other methods. Importantly, all the MCPNet algorithm variants (ℛ^2^, ℛ^3^ and ℛ^4^) and ensemble methods 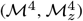 outperformed all existing GRN methods in terms of AUPR.

**Table 4:**
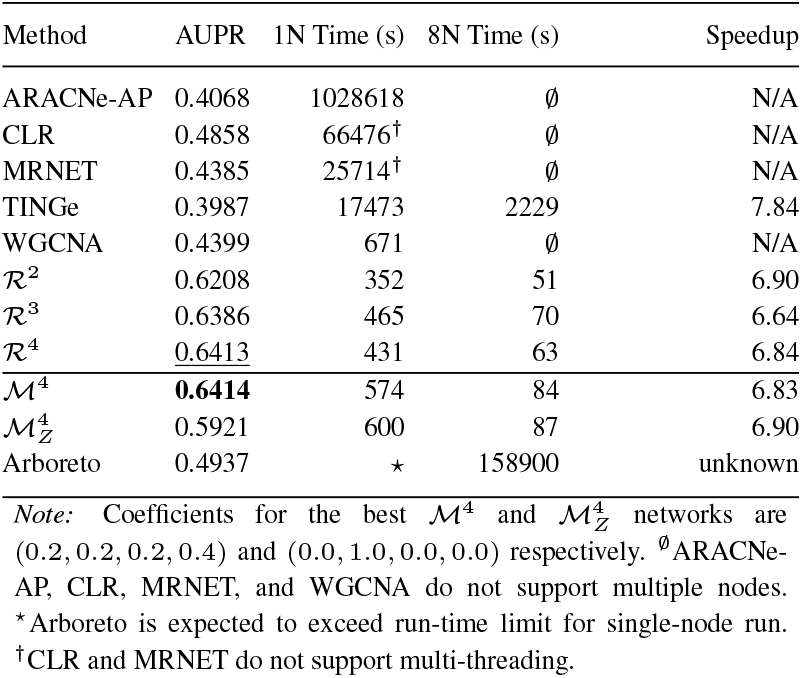
Performance assessment of MCPNet using large-scale *A. thaliana* data. N denotes nodes.

Finally, we evaluate the proposed methods using six *A. thaliana* datasets of different tissues and experimental conditions, each with differing number of samples and genes (see Table 1). Table 5 shows the AUPR and run-times for the multipath ensemble MCP score network (ℳ^4^) and the next best results amongst ARACNe-AP, CLR, MRNET, TINGe, WGCNA, and Arboreto. Complete Results for all the methods are given in supplementary table S3. For all of the datasets analyzed, MCPNet ℳ^4^ shows higher or equivalent AUPR values while the next best AUPR is achieved by CLR, MRNET or Arboreto depending on the datasets.

**Table 5:** AUPR and run-time of ℳ_4_ and the next best results for networks constructed for different *A. thaliana* tissue datasets using 8 24-core nodes. The software that produced the next best AUPR are noted. The first two columns report the input and output matrix sizes as millions of elements.

| Tissue | Mil. Elem. |  | AUPR |  | Run-time (s) |  |
| --- | --- | --- | --- | --- | --- | --- |
|  | Input | Network | MCPNet | 2nd Best | MCPNet | 2nd best |
| Seed | 15.0 | 365.6 | 0.4373 | 0.4285<sup>◇</sup> | 62 | 11298 |
| Flower | 16.2 | 310.0 | 0.5024 | 0.4289<sup>♣</sup> | 52 | 11830 |
| Hormone | 30.3 | 314.5 | 0.4709 | 0.4256<sup>†</sup> | 52 | 26798 |
| Seedling (1wk) | 46.0 | 265.6 | 0.5227 | 0.4424<sup>♣</sup> | 59 | 22336 |
| Leaf | 60.2 | 285.2 | 0.6205 | 0.4529<sup>◇</sup> | 53 | 39202 |
| Development | 93.7 | 337.6 | 0.6414 | 0.4937<sup>†</sup> | 84 | 158900 |

<sup>◇</sup>CLR; <sup>♣</sup>MRNET; <sup>†</sup>Arboreto.

Taken together, our results show that dataset characteristics can significantly influence the quality of the reconstructed network. The coefficients of the weighted average of *L*-path capacities for the best performing ℳ^4^ and 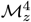 networks also varied by dataset as noted in Tables 3 and 4. The semi-supervised approach therefore is well-suited for the ensemble methods as it adjusts automatically to data characteristics. In addition, since MCPNet is capable of efficiently generating multiple networks in one run (ℛ^2^, ℛ^3^, ℛ^4^, ℳ^4^, and 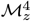), the user can efficiently choose and further evaluate the networks with an appropriately chosen, biologically relevant downstream metric.

### 4.3 Computational Performance of MCPNet

In addition to network quality, Tables 3, 4, and 5 report the run-times and parallel speed up of the proposed and existing methods. The reported times are for the full processing pipelines including file input and output.

Evaluation with the synthetic yeast dataset used a single compute core and 24 shared-memory cores in multi-threaded configuration. Arboreto and Inferelator were run only as multi-threaded as the extrapolated single-core run-time is above job scheduler limit. Table 3 shows that ARACNe-AP, Arboreto and Inferelator respectively required 2.38, 29.07 and 11.17 hours to construct the network using 24-cores, and ARACNe-AP used approximately 29 hours on 1 core to do the same. These represent the worst performing methods.

Of the remaining methods, MCPNet demonstrated the highest absolute computational performance on 24-cores, and on single-core is only bested by WGCNA, which is based on the simpler Pearson correlation and has lower AUPR score than MCPNet. MCPNet is approximately 2× faster than CLR, MRNET, and TINGe using a single core. On 24 cores, TINGe has similar or worse run-time than MCPNet, while CLR and MRNET do not support multi-threading. Interestingly, even though WGCNA supports multi-threading, in practice it experienced a slow-down.

From a parallel scaling perspective, TINGe achieves a nearly perfect speed up of 24× when core count increased from 1 to 24, while our proposed methods obtained speed ups of approximately 17× to 18×. MCPNet’s lower speedups are likely due to the remaining non-parallelizable steps, such as file input and output, occupying a greater proportion of the total run-time.

MCPNet’s performance advantage over the existing methods is more apparent when processing large datasets and in high performance computing environments. Table 4 shows that with one 24-core node, MCPNet outperforms all existing tools – CLR, MRNET, and TINGe required 18.5, 7.1, and 4.9 hours to complete, while at the extreme ARACNe-AP required 285.7 hours. In contrast, MCPNet required between 6 to 10 minutes. The fastest MCPNet method is nearly 3, 000× faster than ARACNe-AP, while the most costly MCPNet method is still more than 1, 700× faster.

The performance advantage of MCPNet is equally striking when 8 nodes (192 cores) are used. Only MCPNet, TINGe, and Arboreto are capable of utilizing multiple nodes. MCPNet significantly outperforms both, up to 43.7× faster than TINGe and 3, 115× faster than Arboreto, when compared using ℛ^2^. The absolute run-times for MCPNet methods ranges between 1 to 1.5 minutes. Compared to the single-node ARACNe-AP run-time, ℛ^2^ is 20, 169× faster. Run-time at this scale renders feasible parameter studies with large gene expression datasets. We also observed parallel scaling factor between 6.64 and 6.90 for MCPNet when node count increased from 1 to 8, while TINGe achieved a closer-to-perfect scaling factor of 7.84 out of 8. TINGe’s longer run-time may be attributed to its higher network communication reliance, while MCPNet’s lower scaling is due to the larger portion of the remaining, unparallelizable components in the pipeline.

Table 5 further illustrates the performance advantage of MCPNet. For the fix datasets with varying sample and gene counts, the ℳ^4^ method, the second most computationally intensive MCP score-based methods, is consistently faster by three to four orders of magnitude when compared to the method that produced the next best network as per AUPR. The (*O*(|*S*||*V*|^2^)) MI computation complexity and the (*O*(|*V*|^3^ log_2_(*L*)) MCP score complexity imply that the run-time of MCPNet scales well with both input sample and gene counts during the MI computation, and with the gene-gene interaction network size during the *L*-path capacity and score computation.

Our evaluations indicate that while dataset characteristics play an important role, the proposed MCPNet methods consistently produce networks that are amongst the highest quality ones based on the AUPR metric. The methods are among the fastest alongside TINGe and WGCNA for smaller datasets on multi-core shared memory machine, and the definitively fastest methods by orders of magnitude for large datasets and in multi-node settings. The combination of network quality and computational performance suggests MCPNet to be a viable first-choice tool for gene regulatory network inference.

## 5 Conclusion

In this paper, we present a new method MCPNet for GRN inference that utilizes a novel edge scoring metric, the MCP score, based on the maximum capacity path (MCP) problem in network and graph analysis. MCP score characterizes long range indirect gene-gene interactions to identify significant direct interactions in an unsupervised manner and with minimal user-specified parameters. We further present a semi-supervised ensemble approach that uses partial network ground truth to optimize the weighted average of MCP scores from multiple *L*-paths scores, with the objective of inferring higher quality novel interactions. The MCP score methods have been shown to generate networks with competitive or superior AUPR scores on real-world *S. cerevisiae* and *A. athaliana* datasets, while reducing run-time by several orders of magnitude relative to existing best-in-class methods. Our efficient parallel implementations allow MCPNet to scale to hundreds of CPU cores, and analyze data with sample and gene counts that were previously prohibitive. With the running time in tens of seconds and higher quality network reconstruction, MCPNet enables analyses of ever-larger datasets in bulk and single cell sequencing as well as parameter space explorations.

## Acknowledgements

This work is supported in part by the National Science Foundation [CCF-1718479].

## Data Availability

The data underlying this article are available in Zenodo, at *https://doi.org/10.5281/zenodo.6499722*. Simulated yeast datasets were generated using scripts in *https://doi.org/10.5281/zenodo.6499756*. Real Yeast dataset (Castro *et al*., 2019) were retrieved from *https://github.com/simonsfoundation/multitask_inferelator/tree/AMuSR.A.athaliana* datasets were derived from NCBI SRA and the accession numbers are listed in Zenodo.

## S1 Supplementary Material

### S1.1 MCP Score and Data Processing Inequality

Previous works on identification of indirect effects in the inferred gene networks are based on Data Processing Inequality (DPI), a property in information theory. In this section, we discuss how DPI-based evaluation can be modeled as a specialization of the *L*-path MCP score defined in section 2.1.

DPI has been adopted by ARACNe-AP and was shown to scale to tens of thousands of genes with TINGe. In these works, DPI-based algorithms compare the direct interaction (MI) to the indirect interactions through a single intermediary.

#### DPI

Let *v*_*s*_ and *v*_*t*_ be two genes and *v*_*i*_ ∈ *V* be some intermediate gene through which *v*_*s*_ and *v*_*t*_ can indirectly interact, forming a Markov chain for the corresponding random variables *X*_*s*_, *X*_*i*_, and *X*_*t*_: *X*_*s*_ *→ X*_*i*_ *→ X*_*t*_. DPI states that for such a Markov chain, mutual information between the genes satisfy the inequalities *W*_*si*_ *≥ W*_*st*_ and *W*_*it*_ *≥ W*_*st*_. If both the conditions *W*_*si*_ *≥ W*_*st*_ and *W*_*it*_ *≥ W*_*st*_ are met, then the Markov chain is a valid information transmission path with an equal or greater capacity than the direct interaction between (*X*_*s*_, *X*_*t*_). Formally, this is expressed as

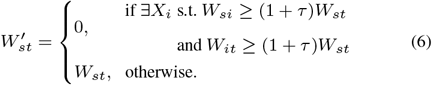

where *W* is the filtered mutual information matrix, and *τ* is a tolerance factor to adjust for data characteristics such as noise. In the cases where 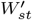 is 0, the direct edge (*v*_*s*_, *v*_*t*_) is considered superfluous and removed.

#### DPI related to MCP Score

The two inequalities in Equation 6 are simultaneously satisfied if the *smaller* of *W*_*si*_ and *W*_*it*_ satisfies the inequality. Furthermore, for 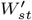 to be set to zero, it suffices that the maximum of min(*W*_*si*_, *W*_*it*_) amongst all possible intermediary genes *v*_*i*_ satisfies the inequality. Equation 6 can therefore be reformulated as below:

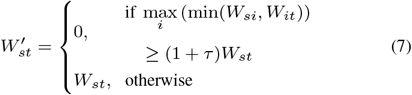

The tolerance parameter *τ* in equations 6 and 7 has to be chosen manually by examining the output and optimized via trial and error. In the absence of ground truth, choosing *τ* is subjective and agnostic of the data. The parameter in Equation 7 can be isolated to suggest a parameter-free gene-gene interaction score *γ*:

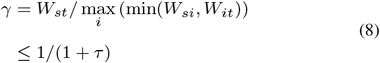

The score *γ* is identical to 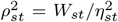 from equations 2 and 3, thus demonstrating that the 2-path MCP score is closely related to DPI.

The 2-path MCP score 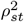 can be interpreted as the relative strength of the direct interactions between genes *v*_*s*_ and *v*_*t*_ to the maximum of all possible indirect interactions between them via some gene *v*_*i*_. In contrast to the DPI-based approach used in ARACNEe and TINGe, 2-path MCP score with MI is directly derived from the gene expression profile data and does not carry discontinuities introduced through a tolerance parameter. Furthermore, MCP scores for all gene pairs also enables precision-recall analyses and better gene interaction predictions based on statistically derived thresholds.

### S1.2 MCPNet Parameter Selection

The MCPNet method has a few parameters: primarily the correlation method used to generate the input for the MCP scoring algorithm and the path length *L*. For the semi-supervised ensemble multipath score, there are two additional parameters: the set of path lengths considered in the ensemble, and the sampling interval during the global search for the optimal *L*-path capacity coefficients. The sampling interval is a user specified parameter that effects a balance between the quality of the global coefficient optimization and compute time constraints. For this paper, we chose to evaluate the method with a sampling interval of 0.1 as the additional compute time this choice incurs is acceptable.

Accurate Estimation of MI from observed data is a difficult problem and numerous methods and multiple surveys have been published (Walters-Williams and Li, 2009; Schaffernicht *et al*., 2010; Doquire *et al*., 2012). While a straightforward method of binning observations and empirically computing MI can be employed (such as by MRNET), the B-spline (Daub *et al*., 2004) and adaptive partitioning (AP; Seok and Kang (2015)) methods are two of the most commonly applied methods in GRN inference. Previous studies also show that these two methods are superior to the binning method(s) in estimating MI (Daub *et al*., 2004; Liang and Wang, 2008). B-spline method, first introduced as an alternative to binning in the context of clustering of gene expression profiles, has been used in many GRN methods including CLR, ARACNe and TINGe. ARACNe-AP (Lachmann *et al*., 2016), the latest update to the ARACNe implementation, uses the AP method for computing MI.

Using simulated yeast data, we first evaluated run-time and accuracy of different MI estimation methods, including our own implementations of B-spline and AP. For B-spline implementations, the typical B-spline parameters of ten (10) bins and three (3) degrees of freedom were employed. Table S1 shows that AP with rank transformed gene expression data as adopted by ARACNe-AP and MCPNet performed better than B-spline based methods with significantly higher AUPR values. AP also has a computational advantage over B-spline based methods. Our implementation based on the same strategy as in Section 2.3 demonstrated superior computational performance for AP when compared to all other B-spline algorithms. Therefore, subsequent studies were all conducted using AP with rank transformation.

We next examined the effect of increasing path length *L* on the quality of the inferred GRN from the simulated Yeast dataset. For each path length *L* between 2 and 30, the *L*-path MCP score is calculated as a score matrix, which is then evaluated against the ground truth network to calculate the AUPR. We observed that AUPR of the MCP scores increases to its maximum rapidly for small *L*, then reduces slightly and settles into a steady oscillation (data not shown). For the simulated yeast dataset and adaptive partitioning MI, the peaks occurs at *L* = 4, and stabilization occurs at *L* = 9. Subsequent evaluations of both MCP scores and the multipath ensemble score employ *L ≤* 4 as a reasonable balance between computational cost and quality improvement.

**Table S1:**
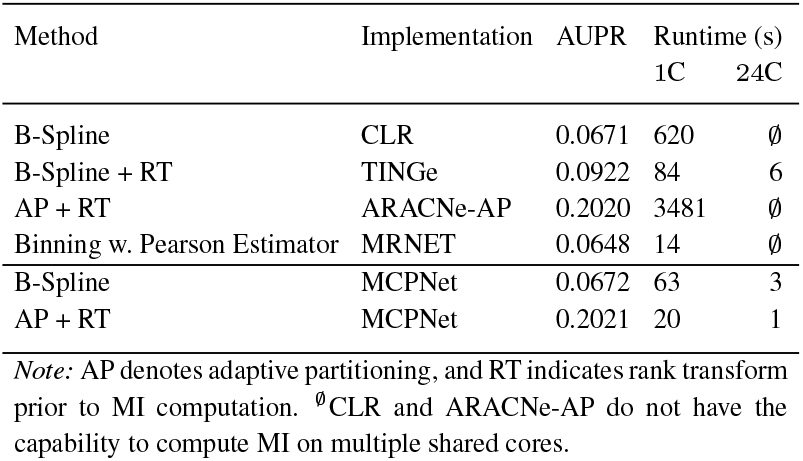
Evaluation of two different MI implementations on simulated yeast data.

## References

Aibar, S. et al. (2017). SCENIC: single-cell regulatory network inference and clustering. Nature Methods, 14(11), 1083–1086.

Aluru, M. et al. (2013). Reverse engineering and analysis of large genome-scale gene networks. Nucleic Acids Research, 41(1), e24–e24.

Aluru, M. et al. (2021). EnGRaiN: a supervised ensemble learning method for recovery of large-scale gene regulatory networks. Bioinformatics.

Bellot, P. et al. (2015). NetBenchmark: a bioconductor package for reproducible benchmarks of gene regulatory network inference. BMC Bioinformatics, 16(1), 1–15.

Bonneau, R. et al. (2006). The Inferelator: an algorithm for learning parsimonious regulatory networks from systems-biology data sets de novo. Genome Biology, 7(5), 1–16.

Brooks, M. D. et al. (2019). Network Walking charts transcriptional dynamics of nitrogen signaling by integrating validated and predicted genome-wide interactions. Nature Communications, 10(1), 1–13.

Castro, D. M. et al. (2019). Multi-study inference of regulatory networks for more accurate models of gene regulation. PLoS computational biology, 15(1), e1006591.

Chockalingam, S. et al. (2016). Microarray data processing techniques for genome-scale network inference from large public repositories. Microarrays, 5(3), 23.

Chockalingam, S. P. et al. (2017). Reverse engineering gene networks: a comparative study at genome-scale. In Proceedings of the 8th ACM-BCB conference, pages 480–490.

Cowen, L. et al. (2017). Network propagation: a universal amplifier of genetic associations. Nature Reviews Genetics, 18(9), 551–562.

Daub, C. O. et al. (2004). Estimating mutual information using b-spline functions–an improved similarity measure for analysing gene expression data. BMC bioinformatics, 5(1), 1–12.

Davis, J. and Goadrich, M. (2006). The relationship between precision-recall and roc curves. In Proceedings of the 23rd ICML, pages 233–240.

Duan, R. and Pettie, S. (2009). Fast algorithms for (max, min)-matrix multiplication and bottleneck shortest paths. In Proceedings of the 20th ACM-SIAM Symposium on Discrete algorithms, pages 384–391.

Faith, J. J. et al. (2007). Large-scale mapping and validation of Escherichia coli transcriptional regulation from a compendium of expression profiles. PLoS Biology, 5(1), e8.

Fernandez, E. et al. (1998). Mosaicking of Aerial Photographic Maps Via Seams Defined by Bottleneck Shortest Paths. Operations Research, 46(3), 293–304.

Hartemink, A. J. (2005). Reverse engineering gene regulatory networks. Nature Biotechnology, 23(5), 554–555.

Huynh-Thu, V. A. et al. (2010). Inferring regulatory networks from expression data using tree-based methods. PloS One, 5(9), 1–10.

Itzhack, R. et al. (2013). Long loops of information flow in genetic networks highlight an inherent directionality. Systems Biomedicine, 1(1), 47–54.

Jin, J. et al. (2015). An Arabidopsis Transcriptional Regulatory Map reveals distinct functional and evolutionary features of novel transcription factors. Molecular Biology and Evolution, 32(7), 1767– 1773.

Lachmann, A. et al. (2016). ARACNe-AP: gene network reverse engineering through adaptive partitioning inference of mutual information. Bioinformatics, 32(14), 2233–2235.

Langfelder, P. and Horvath, S. (2008). WGCNA: an R package for weighted correlation network analysis. BMC Bioinformatics, 9(1), 1–13.

Liang, K.-C. and Wang, X. (2008). Gene regulatory network reconstruction using conditional mutual information. EURASIP Journal on Bioinformatics and Systems Biology, 2008, 1–14.

Lu, L. J. et al. (2007). Comparing classical pathways and modern networks: towards the development of an edge ontology. Trends in biochemical sciences, 32(7), 320–331.

Marbach, D. et al. (2012). Wisdom of crowds for robust gene network inference. Nature Methods, 9(8), 796–804.

Meyer, P. E. et al. (2008). minet: AR/Bioconductor package for inferring large transcriptional networks using mutual information. BMC Bioinformatics, 9(1), 1–10.

Moerman, T. et al. (2019). Grnboost2 and arboreto: efficient and scalable inference of gene regulatory networks. Bioinformatics, 35(12), 2159– 2161.

Pollack, M. (1960). Letter to the Editor—The Maximum Capacity Through a Network. Operations Research, 8(5), 733–736.

Tchourine, K. et al. (2018). Condition-specific modeling of biophysical parameters advances inference of regulatory networks. Cell reports, 23(2), 376–388.

Ullah, E. et al. (2009). An algorithm for identifying dominant-edge metabolic pathways. In 2009 IEEE/ACM International Conference on Computer-Aided Design - Digest of Technical Papers, pages 144–150. ISSN: 1558-2434.

Vassilevska, V. et al. (2007). All-pairs bottleneck paths for general graphs in truly sub-cubic time. In Proceedings of the thirty-ninth annual ACM symposium on Theory of computing, STOC ‘07, pages 585–589, New York, NY, USA. Association for Computing Machinery.

Vermeirssen, V. et al. (2014). Arabidopsis ensemble reverse-engineered gene regulatory network discloses interconnected transcription factors in oxidative stress. The Plant Cell, 26(12), 4656–4679.

Zola, J. et al. (2010). Parallel information-theory-based construction of genome-wide gene regulatory networks. IEEE Transactions on Parallel and Distributed Systems, 21(12), 1721–1733.

## SI References

Doquire, G. et al. (2012). A comparison of multivariate mutual information estimators for feature selection. In ICPRAM (1), pages 176–185.

Schaffernicht, E. et al. (2010). On estimating mutual information for feature selection. In International Conference on Artificial Neural Networks, pages 362–367. Springer.

Seok, J. and Kang, Y. S. (2015). Mutual information between discrete variables with many categories using recursive adaptive partitioning. Scientific reports, 5(1), 1–10.

Walters-Williams, J. and Li, Y. (2009). Estimation of mutual information: A survey. In International Conference on Rough Sets and Knowledge Technology, pages 389–396. Springer.

